# Veterinary insecticides in wild bird nests: emerging contaminants in urban and protected forest habitats

**DOI:** 10.64898/2026.06.10.731177

**Authors:** Zsófia Varga-Szilay, Olivér Csíkvári, Olga Ulbert, Ivett Pipoly, Gábor Seress

**Author notes:** Corresponding author: Zsófia Varga-Szilay.

## Abstract

Veterinary ectoparasiticides widely used on companion animals often contain synthetic insecticides whose agricultural applications have been restricted in the European Union because of environmental concerns. These neurotoxic compounds can persist on animal fur and enter surrounding environments, potentially exposing non-target organisms. The Great Tit (*Parus major*), a common passerine in both urban and forest habitats, frequently incorporates animal fur in nest linings, thereby creating a potential exposure route for breeding adults and nestlings. We analysed fur-containing nest material collected from 63 Great Tit nests in artificial nest boxes at an urban site and a nearby protected forest in Hungary during the 2025 breeding season, sampling at mid-nestling stage and also after fledging. Using HPLC-MS/MS and GC-MS, we detected several veterinary ectoparasiticides in nest materials, including fipronil, fipronil sulfone, imidacloprid, and permethrin. Acetamiprid was found only in urban nests, indicating additional, non-veterinary environmental sources. Multiple insecticides were present in nest material, with higher contamination levels and greater compound diversity in urban compared to forest nests. Residues were present at both sampling times but declined over the course of the breeding cycle. Although contamination was not associated with the measured reproductive parameters of Great Tits, our findings show that veterinary ectoparasiticides can contaminate wild bird nests, including those in protected forest ecosystems. This highlights a previously under-recognised pathway linking companion animal treatments to wildlife exposure and underscores the need to assess the ecological risks and trade-offs associated with widespread veterinary insecticide use.

**Highlights:**

- Veterinary ectoparasiticides were detected in urban and forest Great Tit nests
- Urban nests contained higher contamination levels and greater compound diversity
- Acetamiprid occurred only in urban nests, indicating additional environmental inputs
- Protected forests also exposed to fipronil, fipronil sulfone and permethrin
- Bird nests reveal an overlooked pathway linking pet treatments to wildlife exposure

## 1. Introduction

Synthetic pesticides are among the most pervasive chemical stressors in human-altered ecosystems (Wan et al., 2025). Increasing evidence demonstrates that their persistence, mobility, and bioaccumulation can generate long-term and cascading ecological effects on non-target organisms, disrupting trophic interactions (Hallmann et al., 2014; Molenaar et al., 2024; Montaigu & Goulson, 2020).

Veterinary ectoparasiticides used in livestock and companion animal treatments consist predominantly of three major insecticide classes: neonicotinoids (e.g. imidacloprid), phenylpyrazoles (e.g. fipronil), and synthetic pyrethroids (e.g. permethrin). These compounds act by disrupting neural signalling in arthropods, leading to paralysis and mortality of target parasites (Li & Akk, 2008; Tomizawa & Casida, 2005). However, growing evidence indicates that they may also affect non-target organisms through direct toxicity, sublethal physiological and behavioural effects, and indirect ecological pathways such as prey depletion (Goulson, 2013; Pisa et al., 2021).

Although some of these active ingredients are no longer approved or are severely restricted for agricultural use in the European Union due to their risks to pollinators and other non-target organisms (Goulson & 232 Signatories, 2018; Joachim et al., 2025), they remain widely used in veterinary contexts (Pesticide Action Network UK, 2023). Despite this continued application, relatively little is known about their environmental dissemination and potential exposure routes in terrestrial ecosystems, particularly in relation to avian breeding habitats.

Following topical administration, ectoparasiticides can persist on treated animals’ fur for extended periods and may be transferred to the environment through shedding, grooming, and contact with surfaces (Preston-Allen et al., 2023). As a result, these compounds have been detected in a range of environmental matrices, including gardens, urban green spaces, agricultural soils, and aquatic systems (Perkins et al., 2025; Perkins & Goulson, 2023), raising concerns about their broader ecological consequences (Pisa et al., 2021).

Bird nests may represent a previously under-recognised exposure pathway. Many bird species incorporate animal fur into nest linings (Hansell, 2000; Harničárová & Adamík, 2016) thereby potentially introducing veterinary insecticides directly into avian breeding environments. This pathway may be particularly important in urban habitats where, owing to the scarcity of large wild mammals, fur used as nesting material frequently originates from domestic animals (Girão et al., 2024). However, contamination may also extend into forest habitats because dogs accompanying hikers and hunters are commonly treated with veterinary ectoparasiticides and may introduce contaminated fur and residues into natural or semi-natural environments. Consequently, residues of veterinary ectoparasiticides transferred in this manner may result in direct exposure of breeding adults, eggs, and developing nestlings.

The Great Tit (*Parus major*) is a small-bodied, insectivorous passerine widely distributed across Europe, occupying habitats ranging from natural woodlands to highly urbanised areas. The species is a facultative double-breeder, meaning that part of the pairs lay and raise two clutches within a breeding season. Great Tits readily use artificial nest boxes and frequently incorporate animal fur into their nests (Lambrechts et al., 2017; Mainwaring et al., 2012), making it a suitable model for assessing contaminant transfer and ecological consequences in wild bird populations. Empirical evidence supports the plausibility of this exposure route; for example, Guldemond et al. (2019) detected 26 different insecticides, including fipronil and imidacloprid, in deceased Great Tit nestlings. More recently, pesticide residues have been reported in all examined nest materials of Great Tits and Blue Tits (*Cyanistes caeruleus*), breeding in residential gardens, with high prevalence associated with increased chick mortality (Tassin-de-Montaigu et al., 2025). These findings suggest that veterinary insecticides may represent an overlooked risk factor affecting reproductive success in urban and peri-urban bird populations.

On the other hand, exposure to insecticides within nests may also involve ecological trade-offs. Synthetic pyrethroids and other ectoparasiticides can reduce the abundance of nest-dwelling arthropod parasites, potentially improving nestling condition and survival under high parasite pressure. For example, permethrin treatments have been shown to reduce ectoparasite loads, as Bulgarella et al. (2020) concluded that under certain conditions, fitness benefits may outweigh potential sublethal physiological effects. Similarly, insecticide-based control of the invasive nest fly *Philornis downsi* has been shown to improve reproductive success, with permethrin treatments significantly increasing fledging success in Darwin’s finches, although outcomes varied with concentration and application method (Kofler et al., 2025). Comparable dual effects have been documented in urban bird nests containing cigarette butts, which reduce parasite loads while simultaneously imposing oxidative and genotoxic stress on chicks (Suárez-Rodríguez et al., 2013; Suárez-Rodríguez & Macías Garcia, 2014). Thus, the net fitness consequences of veterinary insecticide contamination in bird nests remain uncertain and are likely determined by compound identity, concentration, exposure duration, and parasite pressure.

In this study, we evaluate the extent of veterinary insecticide contamination in Great Tit nest materials and examine its potential consequences for reproductive performance. By including both urban and forest populations, we assess whether contamination and its potential biological consequences extend beyond urban environments into nearby habitats that are generally considered less affected by human activities. Specifically, we quantify insecticide concentrations in fur-containing nest materials and compare contamination levels between urban and forest habitats, assess temporal changes in contamination within nests during both the breeding season and the breeding cycle, and evaluate association between contamination and reproductive parameters, including the proportion of unhatched eggs and nestling mortality within nests, as well as body size of fledglings.

We hypothesise that (**H1**) urban nests will contain a greater diversity and higher concentrations of insecticides than forest nests; and (**H2**) insecticide contamination will be lower during the seasonally first than during the second brood period, while insecticide concentrations will decrease over time within individual nests. Furthermore, (**H3**) higher insecticide contamination will be associated with lower rates of hatching success, higher rates of nestling mortality, as well as reduced nestling body size prior to fledging. By elucidating this exposure pathway, our study contributes to a better understanding of the environmental occurrence and spread of veterinary insecticides in terrestrial habitats and their potential implications for wild bird populations.

## 2. Material and Methods

### 2.1 Study framework and sites

The fieldwork and data collection were conducted within the framework of a long-term, ongoing project (HUN-REN Hungarian Research Network and the National Research, Development and Innovation Office, FK-137743). The urban study site is located in Veszprém, a medium-sized city in Hungary (47°05′17.29″N, 17°54′29.66″E), where nest boxes are distributed across public green spaces spanning the urbanisation gradient. The forest study site is located near the village of Farkasgyepű in a protected Natura 2000 forest (47°11’48’N, 17°39’35’E) dominated by European beech (*Fagus sylvatica*) and hornbeam (*Carpinus betulus*), located ca. 30 km from Veszprém (**Supplementary Material Figure 1**).

Nests are checked at least twice per week throughout the breeding season, from egg laying to fledging, covering both the annual first and second breeding (March–July) attempts of Great Tits. During these checks, breeding parameters, like the number of eggs, dates of incubation initiation and hatching, number of hatchlings, and number of fledglings are recorded. Before fledging, nestlings are measured and ringed at an age of 14 to 16 days post-hatch (taking the day of hatching as day 1). For each nestling, we recorded body mass (±0.1 g) and measured the length of the left tarsus (±0.1 mm) and right wing (from the bend of the wing to the tip of the longest primary, following Svensson’s ‘alternative’ method (Svensson, 1992); ±1 mm).

### 2.2 Sample collection and preparation

During the breeding season of 2025, nest material containing animal fur was collected at two time points from the sampled nests: first, when nestlings were 4-6 days old (mid-term sampling; MT), and second, within a few days after fledging (final sampling; F). In the mid-term sampling, a small subsample of nest material containing animal fur was carefully removed using metal forceps and placed into labelled, sealable plastic zip-lock bags. To prevent cross-contamination, forceps were thoroughly wiped with 70% ethanol before and after handling each sample. During the final sampling, the entire nest material was collected from the nest box and stored individually in labelled plastic bags. In the laboratory, all samples were stored at −20°C to prevent degradation of insecticide residues. Prior to chemical analysis, animal fur was manually separated from the remaining nest material using a stereomicroscope and an analytical precision balance. Where sufficient material was available, up to 25 mg of animal fur per nest was subsampled. Eight samples contained less than 25 mg of fur; however all provided sufficient material for chemical analysis. The subsamples were transferred into sterile 15 mL polypropylene centrifuge tubes with screw caps, and individually labelled and transported in these tubes to the analytical laboratory for further preparation and insecticide residue analysis. In total, 63 nest-lining material samples were obtained from 40 broods (n = 23 broods with MT and F samples, n = 4 broods with MT samples only, and n = 13 broods with F samples only).

### 2.3 Chemical analysis

We analysed seven synthetic insecticides representing the principal classes of veterinary ectoparasiticides used in companion animals, including the neonicotinoids imidacloprid, dinotefuran, and nitenpyram (primarily used in oral veterinary formulations), the phenylpyrazole fipronil and its major degradation product fipronil sulfone, and the synthetic pyrethroid permethrin. Imidacloprid, fipronil, and permethrin are among the most widely used active ingredients in topical veterinary ectoparasite treatments for companion animals (Wells & Collins, 2022), whereas acetamiprid was included because of its widespread use as a systemic neonicotinoid insecticide.

Blank fur samples collected from dogs known to be untreated with the compounds of interest were collected and used as a matrix for blank, calibrator and QC samples.

### 2.4 Sample extraction

25 mg fur subsamples were spiked with 5 µL of 10µM Glyburide solution in acetonitrile. For extraction, 1000 µL of HPLC grade acetonitrile was added to each sample, vortexed for 10 seconds, then samples were placed in an ultrasonic bath for 15 minutes. After sonication, samples were vortexed again, then extracts were transferred into 5 mL Eppendorf tubes. Acetonitrile extraction was repeated one more time, followed by a third extraction with 1500 µL of acetonitrile:water (2:1). All extracts were combined. 1000 µL of brine was added to the extracts, and the tubes were vortexed for 30 seconds. After phase separation, 1800 µL of the supernatant was transferred into QuEChERS Clean-up tubes containing 150 mg MgSO4, 25 mg PSA, 25 mg C18 and vortexed for 30 seconds. All samples were centrifugated at 5°C, 3000 RPM for 10 minutes. After centrifugation, two 500 µL aliquots of the supernatant were transferred into 2 mL polypropylene deep well plates and evaporated to dryness under nitrogen. For LC-MS/MS analysis one batch of the samples was reconstituted in 150 µL acetonitrile:water (1:1), the other batch in 150 µL methanol for GC-MS analysis. Samples were stored at -20°C until analysis.

For calibrator and QC samples, blank fur samples were spiked with 8 µL aliquots of a dilution series in acetonitrile containing dinotefuran, fipronil, fipronil sulfone, imidacloprid, and nitenpyram, and went through the same extraction procedure. Blank samples were spiked with 13 µL acetonitrile before extraction and were prepared without adding internal standard.

### 2.5 Sample analysis

Insecticide content was analysed using HPLC-MS/MS (dinotefuran, fipronil, fipronil sulfone, imidacloprid, and nitenpyram) and GC-MS (permethrin).

#### 2.5.1. LC-MS/MS analysis

Analysis of dinotefuran, fipronil, fipronil sulfone, imidacloprid, and nitenpyram was performed on an LS-I autosampler (Sound Analytics, Niantic, CT, USA) equipped with Agilent 1260 HPLC pumps coupled to a Sciex 6500+ Triple Quadrupole Mass Spectrometer (AB SCIEX, Framingham, MA, USA). Chromatographic separation was achieved on a Phenomenex Kinetex F5 column (30 × 2.1 mm, 2.6 µm, Phenomenex Inc., Torrance, CA, USA) in 1.4 minutes with a gradient elution described in **Supplementary Material Table 1** Eluent A consisted of 5 mM ammonium-acetate and 1 mM ammonium-fluoride in water, eluent B consisted of 5 mM ammonium-acetate and 1 mM ammonium-fluoride in acetonitrile:water (8:2). Mass spectrometric detection was performed in Multiple Reaction Monitoring mode, with the MRM parameters optimized for each analyte during method development (**Supplementary Material Table 2**). Lower limits of quantitation (LLOQ) for each analyte are shown in **Supplementary Material Table 3** Chromatograms were evaluated using Analyst software (AB SCIEX, Framingham, MA, USA).

#### 2.5.2. GC-MS analysis

Permethrin concentration in the extracts prepared according to section 2.4 was determined on a Shimadzu GCMS-QP2010 SE instrument (Shimadzu Corporation, Kyoto, Japan) equipped with split/splitless injector. Chromatographic separation was achieved on a ZB-5MS plus column (30 m, 0.25 mm, 0.25 µm) (Phenomenex, Torrance, USA) applying an oven temperature program between 60-310°C. The total analysis time was 15.50 min. A hybrid scan/SIM MS method was applied allowing for the detection of matrix components in scan mode and the sensitive measurement of permethrin in SIM mode. Details of the GC-MS method and the MS detection parameters are presented in **Supplementary Material Table 4**.

### 2.6 Statistical analysis

For insecticide contamination, we used two complementary metrics: first the total number of detected insecticides, representing the number of insecticides (including permethrin); and second, the total insecticide concentration (excluding permethrin because only its presence or absence could be determined) detected in the nest sample.

To compare contamination levels between habitats (**H1**), we analysed differences in both the total number of detected insecticides and the total insecticide concentration between urban (n = 43) and forest (n = 20) samples (from n = 29 urban and n = 11 forest broods). The total number of insecticides was analysed using a generalised linear mixed-effects model (GLMM) with Poisson error distribution. Total insecticide concentration was log-transformed and analysed using a linear mixed-effects model (LMM). In both models, habitat (urban or forest) was included as fixed effects, and nest identity was included as a random intercept.

To assess temporal changes in insecticide contamination in nests, we restricted these analyses to the urban habitat only, where we expected the nest material to be more strongly exposed to chemical contamination (**H2a**). To do so, we compared both the total number of detected insecticides and their total concentration between samples collected during the seasonally first (n = 28 samples) and second (n = 15 samples) brood periods during the breeding season (total: n = 29 broods). The total number of insecticides was analysed using a generalised linear mixed-effects model (GLMM) with a Conway–Maxwell–Poisson error distribution to account for dispersion. Residual diagnostics indicated significant underdispersion (dispersion = 0.537, p = 0.032) supporting the choice of this distribution. Total insecticide concentration was log-transformed and analysed using a linear mixed-effects model (LMM). In both models, brood type (first or second brood) was included as a fixed effect, and nest identity was included as a random intercept. Moreover, to assess temporal changes in total insecticide concentration within a breeding event in urban nests ( **H2b**), we analysed the total insecticide concentration between mid-term (n = 14) and final (n = 14) samples (total: n = 14 broods). This was analysed using a linear mixed-effects model (LMM) with log-transformed concentration as the response variable, sample type (MT or F) as a fixed effect, while nest identity was included as a random intercept.

To evaluate associations between insecticide contamination and Great Tits’ reproductive parameters, we involved urban nests only. To calculate hatching success, we used the proportion of unhatched eggs and clutch size (number of eggs counted during the incubation period); to calculate nestling mortality, we used the proportion of dead chicks and number of hatched chicks. Eggs of unknown fate were assumed to have hatched but the chicks died shortly afterwards and were subsequently removed from the nest by the parents (as early-age nestling mortality is typical in urban Great Tits; pers. obs.; Corsini et al., 2021). In addition, all chicks that were found dead in the nest, as well as chicks known to have hatched but missing at the time of inspection, were classified as dead chicks. Finally, we excluded all nests where complete brood loss was probably attributable to nest predation (evidenced by missing brood, predator marks on dead chicks or damage to the nest structure) either during the egg stage (in case of hatching success models) or during the nestling-rearing period (in case of nestling mortality models). Hatching success (**H3a**; n = 27) and nestling mortality (**H3b**; n = 24) were analysed in generalised linear mixed-effects models (GLMMs) with a beta-binomial error distribution. Response variables were specified as the counts of unhatched versus hatched eggs, and dead versus surviving chicks, respectively. Separate models were fitted including either the highest number of total insecticides detected in the sample or the highest detected total insecticide concentration (in nests where both MT and F samples were available) as predictor variables, together with brood type (first or second) as a fixed effect.

Nestling (n = 103) morphological traits (**H3c**; tarsus length, wing length, and body mass) were analysed using linear mixed-effects models (LMMs) (n = 21 urban broods). Separate models were fitted for each response variable, including either the highest insecticide number detected in the sample or the highest total insecticide concentration as predictor variables. Brood type (first or second) was included as a fixed effect, and nest identity as a random intercept.

The data analysis and visualisation were conducted in the R software environment (R Core Team, 2021). Mixed-effects models were fitted using the *lme4* (Bates et al., 2015) and *glmmTMB* (Brooks et al., 2017) packages. Linear mixed-effects models were evaluated through visual inspection of residuals and Q–Q plots. For generalised linear mixed-effects models, residual uniformity and dispersion were assessed using the *DHARMa* package (Hartig et al., 2024). Figures were created using *ggplot2* (Wickham, 2016).

Note that nest-box identity was not included in our models as a random factor (to control for the potential non-independence of the data resulting, for example, from territory quality) as we had only two cases in our dataset when a seasonally first and second brood was initiated in the same nest box.

## 3. Results

### 3.1 Insecticide contamination in urban and forest nests

Out of 63 fur samples from Great Tit nests, 31.75% (n = 20) were from forests and 68.25% (n = 43) were from urban habitats. Urban nests were contaminated with fipronil, fipronil sulfone, imidacloprid, and permethrin, whereas forest nests contained fipronil sulfone, imidacloprid, and permethrin. In addition, acetamiprid was also detected in the nest material in 8 of the 43 urban samples (**Table 1**).

**Table 1:** Insecticides detected in Great Tit nests are presented together with the number of samples (n), the percentage of samples exceeding the Lower Limit of Quantification (% > LLOQ); for permethrin, LLOQ 74.14 ppb), as well as the maximum (max CO), median (median CO), and mean concentration (average CO ± SE), expressed in part per billion (ppb).

| Compound | Forest (n = 20) |  |  |  |  | Urban (n = 43) |  |  |  |  |
| --- | --- | --- | --- | --- | --- | --- | --- | --- | --- | --- |
| | n | % > LLOQ | Max CO | Median CO | Average CO (±SE) | n | % > LLOQ | Max CO | Median CO | Average CO (±SE) |
| Acetamiprid | 0 | 0 | - | - | - | 8 | 18.60 | 225 | 22.32 | 45.05 (25.84) |
| Dinotefuran | 0 | 0 | - | - | - | 0 | 0 | - | - | - |
| Fipronil | 0 | 0 | - | - | - | 27 | 32.79 | 579.24 | 21.27 | 72.16 (23.92) |
| Fipronil sulfone | 4 | 20.00 | 8.18 | 3.96 | 4.75 (1.15) | 35 | 81.40 | 459 | 13.32 | 45.82 (17.23) |
| Imidacloprid | 2 | 10.00 | 15.12 | 15.12 | 10.8 (4.32) | 8 | 18.60 | 674.28 | 64.44 | 137.25 (77.44) |
| Nitenpyram | 0 | 0 | - | - | - | 0 | 0 | - | - | - |
| Permethrin | 1 | 5.00 | < LLOQ | < LLOQ | < LLOQ | 13 | 30.23 | < LLOQ | < LLOQ | < LLOQ |

We found that both insecticide number (GLM, β = 1.800 ± 0.392, p < 0.001; **Figure 1A**) and concentration (LMM, β = 2.787 ± 0.527, p < 0.001; **Figure 1B**) were significantly higher in urban than forest nests. The highest concentration of imidacloprid was detected in an urban sample (674.28 ppb); moreover, the highest total insecticide concentration measured in a nest was 946.1 ppb. The total concentration of insecticides ranged from 3.24 ppb to 674.28 ppb, with an average of 60.13 ppb (13.15±SE).

**Figure 1:**
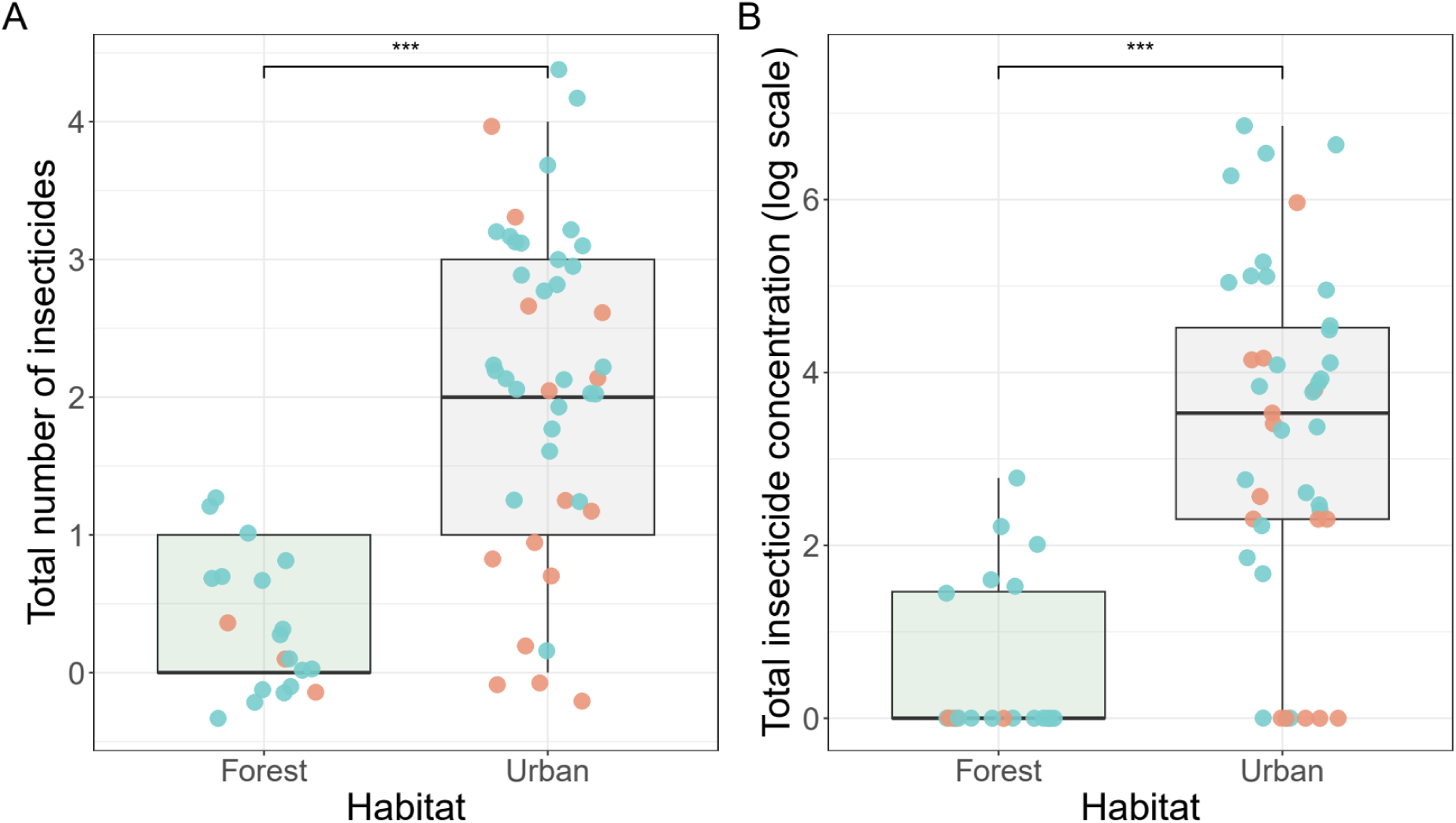
(A) Total number of detected insecticides and (B) total insecticide concentration (log-transformed) in nest materials of Great Tits from forest (n = 20) and urban (n = 43) habitats. Forest and urban habitats are represented by light green and grey boxplots, respectively, showing the median, interquartile range, and data distribution. Asterisks indicate the statistical significance of differences between habitats (* p < 0.05, ** p < 0.01, *** p < 0.001). Blue dots represent the seasonally first, red dots the second broods.

### 3.2 Temporal changes in insecticide contamination during the breeding cycle in urban nests

The number of detected insecticides changed significantly during the breeding season. The total number of insecticides was significantly lower in second broods compared to first broods (GLMM, β = −0.519 ± 0.192 SE, p = 0.007; **Figure 2A**). Similarly, the total insecticide concentrations were significantly lower in second broods compared to first broods (LMM, β = −1.457 ± 0.669 SE, p = 0.040) (**Figure 2B**).

**Figure 2:**
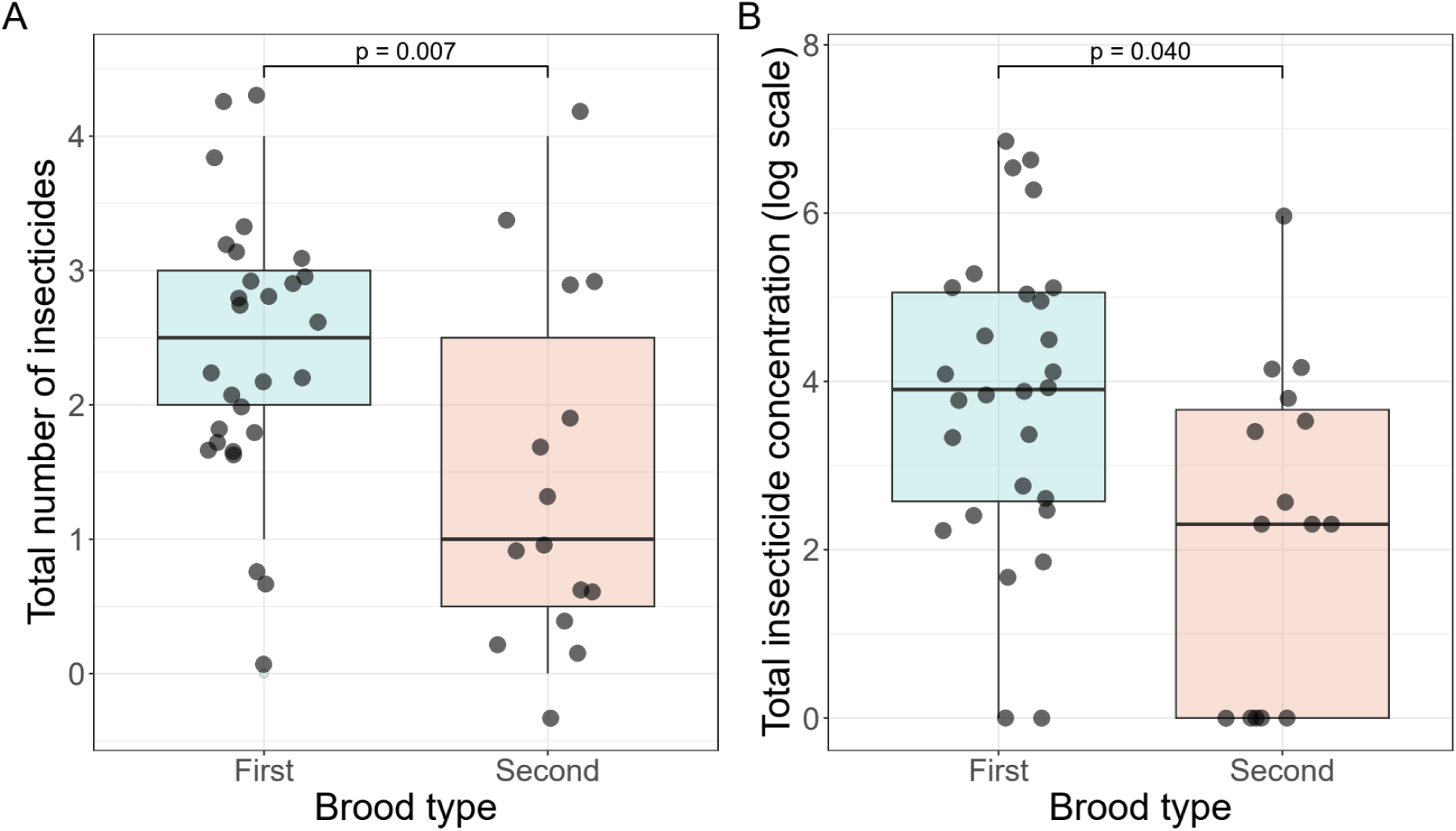
(A) Total number of detected insecticides and (B) total insecticide concentration (log-transformed) in nest materials of urban Great Tits from first (n = 28) and second (n = 15) broods. Boxplots show medians and interquartile ranges, with points representing individual nests. Statistical differences between brood types are indicated in each panel with exact p-values.

Within-nest temporal variation revealed a significant difference between mid-term and final samples (**Figure 3**). Total insecticide concentrations (log-transformed) were significantly higher in mid-term samples compared to final samples (LMM, β = 1.252 ± 0.430 SE, p = 0.012).

**Figure 3:**
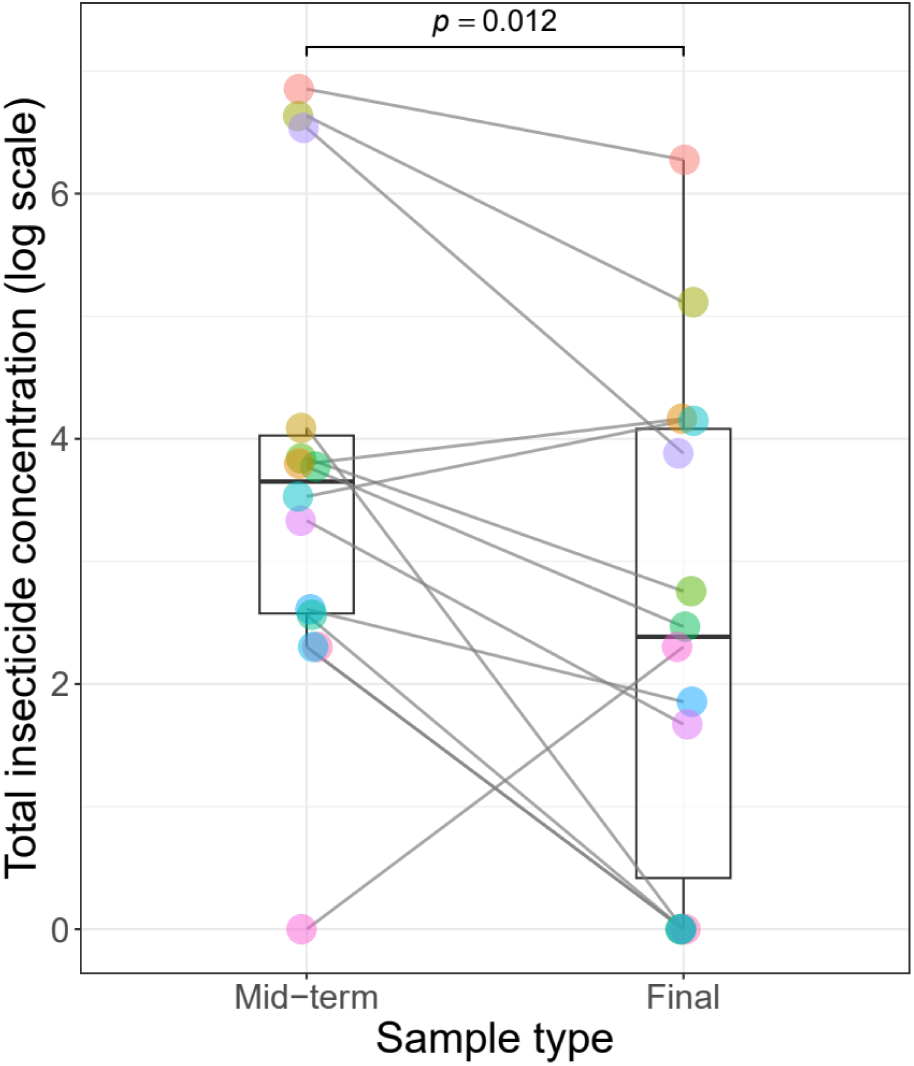
Total insecticide concentration (log-transformed) measured in mid-term and final samples from urban Great Tit nests during the breeding period. Boxplots represent the median, interquartile range, and overall distribution of values within each sampling period. Dots are coloured by nest identity, with each colour representing an individual nest (n = 14) that has both mid-term and final samples. Connecting lines link paired samples from the same nest, illustrating within-nest temporal changes between sampling periods.

### 3.3 Associations between insecticide contamination and reproductive parameters in urban nests

#### Hatching success

We found no significant relationship between insecticide contamination and the proportion of unhatched eggs, either for the total number of insecticides (GLMM, β = 0.092 ± 0.434, p = 0.833) or for total insecticide concentration (GLMM, β = −0.007 ± 0.260, p = 0.980). In addition, brood type (first or second brood) had no significant effect on hatching success in either model (p = 0.435 and p = 0.391, respectively).

#### Nestling mortality

Similarly, we observed no significant relationship between insecticide contamination and nestling mortality, either for the total number of insecticides (GLMM, β = -0.084 ± 0.200, p = 0.676) or for total insecticide concentration (GLMM, β = 0.007 ± 0.125, p = 0.957) (**Figure 4**). However, brood type significantly affected offspring survival, with seasonally second broods having higher offspring mortality than first broods in both models (β = -1.354 ± 0.504, p = 0.007; and β = 1.437 ± 0.501 p = 0.004, respectively).

**Figure 4:**
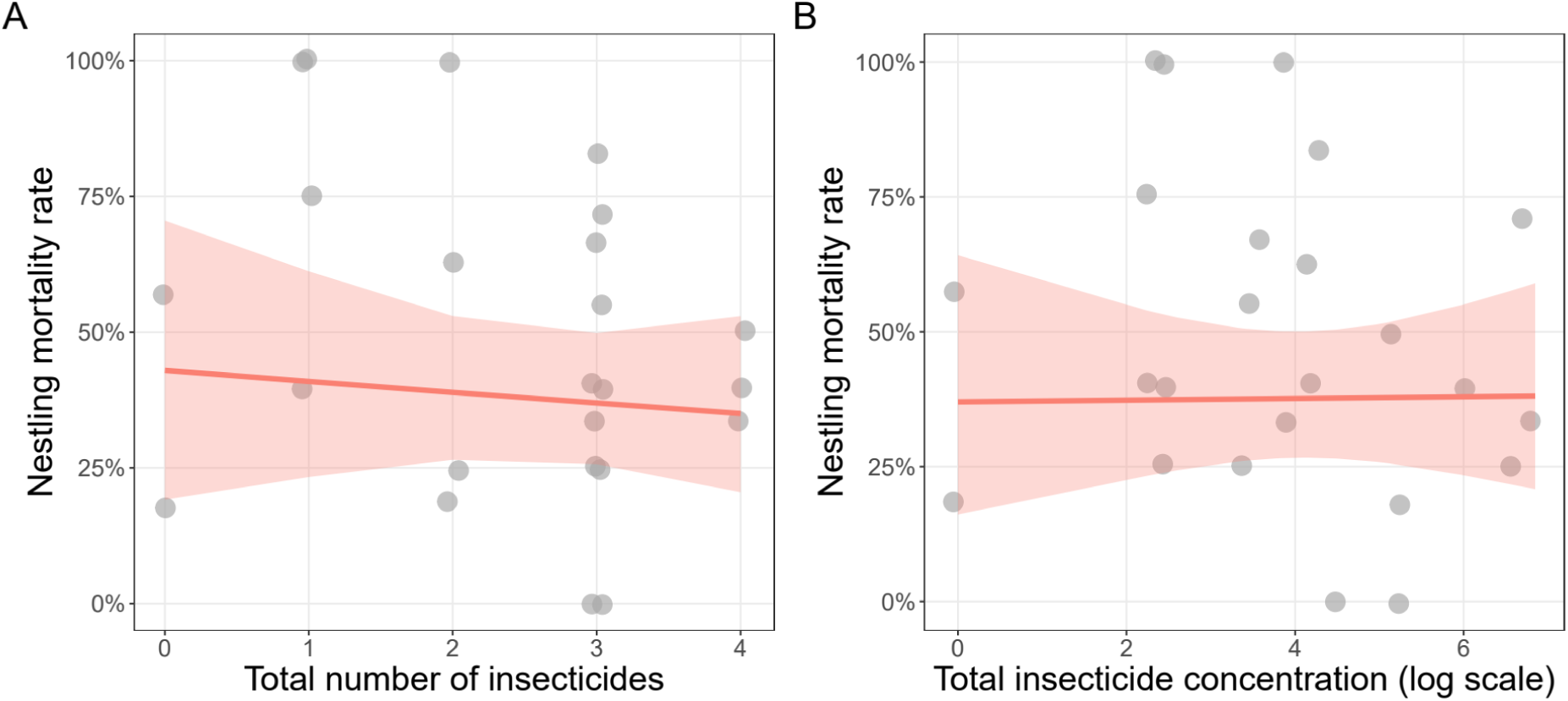
Relationship between nestling mortality rate and insecticide exposure. (A) Mortality rate as a function of the number of detected insecticides. (B) Mortality rate as a function of total insecticide concentration (log-transformed). Gray dots represent broods (n = 24). Solid red lines show predictions from beta-binomial generalised linear models, with shaded areas indicating 95% confidence intervals.

#### Nestling body size parameters

Nestling body size parameters, including tarsus length, wing length, and body mass, were not significantly associated with either the highest number of detected insecticides or the highest total insecticide concentration (**Table 2**, **Figure 5**). However, brood type significantly affected tarsus length, with nestlings from second broods having shorter tarsi compared to those from first broods (**Table 2**). No significant effect of brood type was detected for wing length or body mass.

**Table 2:** Associations between insecticide contamination and nestling (n = 103) body size parameters (tarsus length, wing length, and body mass) estimated using linear mixed-effects models (LMMs). Insecticide exposure was expressed as either the highest number of detected insecticides or the highest detected total insecticide concentration within the nest-lining material (from n = 21 broods). Significant effects (p < 0.05) are shown in bold.

| Response variable | Predictor | $\beta \pm SE$ | t | p |
| --- | --- | --- | --- | --- |
| Tarsus length<br>(in mm) | Highest insecticide count | $-0.076 \pm 0.143$ | -0.53 | 0.60 |
|  | **Brood type (second)** | **$-0.993 \pm 0.389$** | **-2.56** | **0.02** |
| | Highest insecticide concentration | $-0.109 \pm 0.081$ | -1.34 | 0.20 |
|  | **Brood type (second)** | **$-1.101 \pm 0.361$** | **-3.05** | **0.01** |
| Wing length<br>(in mm) | Highest insecticide count | $-0.225 \pm 0.985$ | -0.23 | 0.82 |
| | Brood type (second) | $-3.278 \pm 2.570$ | -1.28 | 0.22 |
| | Highest insecticide concentration | $-0.459 \pm 0.575$ | -0.8 | 0.44 |
| | Brood type (second) | $-3.843 \pm 2.442$ | -1.57 | 0.13 |
| Body mass<br>(in grams) | Highest insecticide count | $0.205 \pm 0.390$ | 0.53 | 0.61 |
| | Second brood | $-1.120 \pm 1.038$ | -1.08 | 0.29 |
| | Highest insecticide concentration | $-0.147 \pm 0.230$ | -0.64 | 0.53 |
| | Brood type (second) | $-1.656 \pm 0.998$ | -1.66 | 0.11 |

**Figure 5:**
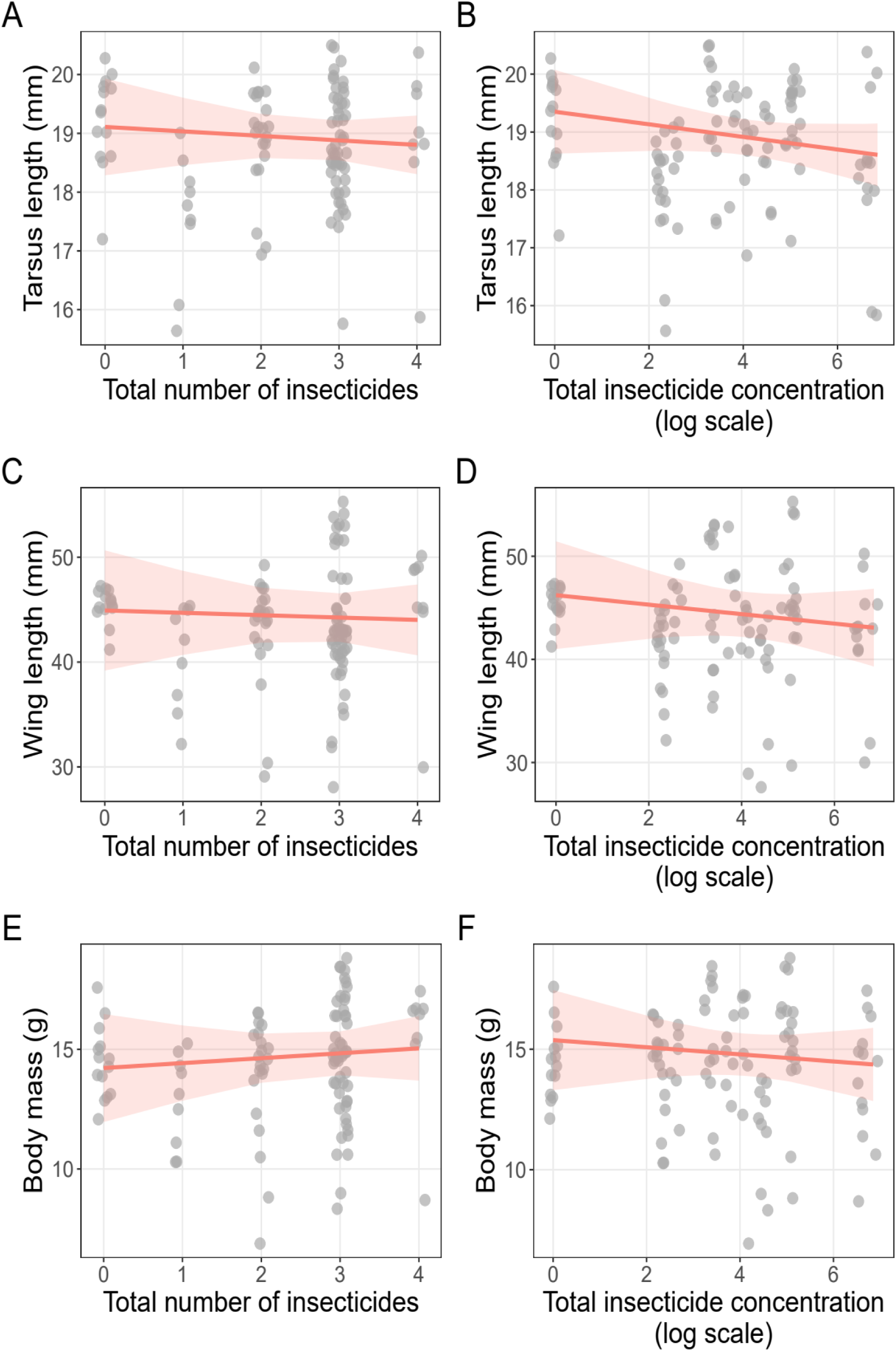
Relationships between nestling body size parameters (A, B: tarsus length; C, D: wing length; and E, F: body mass) and insecticide contamination, expressed as the total number of detected insecticides (left panels) and log-transformed total insecticide concentration (log-transformed; right panels). Red lines represent predictions from linear mixed-effects models (LMMs), while shaded areas indicate 95% confidence intervals. Grey dots represent raw observations (n = 103 offspring measurements from 21 broods). All models included nest identity as a random effect.

## 4. Discussion

Consistent with recent findings (Tassin-de-Montaigu et al., 2025), we detected fipronil, fipronil sulfone, imidacloprid, and permethrin in nest-lining materials, confirming that veterinary insecticides can disperse into surrounding environments and become incorporated into avian nests through the collection of contaminated mammalian fur.

Supporting our first hypothesis (**H1**), urban nests showed both higher diversity and higher concentrations of insecticides compared to forest nests. This pattern likely reflects greater exposure opportunities in urban and peri-urban environments, where companion animals frequently use shared green spaces and routine flea and tick treatments are commonly applied (Jennett et al., 2013; Perkins & Goulson, 2023). Treated pets likely shed contaminated fur continuously across a wide range of urban microhabitats, including parks, domestic gardens, residential green spaces, street verges, and other semi-natural habitats embedded within the urban matrix (Preston-Allen et al., 2023), many of which are regularly used by Great Tits and other insectivorous birds (Lakatos et al., 2025). Consequently, urban habitats may function as accumulation zones where multiple overlapping contamination sources increase the likelihood of insecticide residues being incorporated into nest-lining material.

The predominance of fipronil (a veterinary ectoparasiticide) and its degradation product, fipronil sulfone, in urban nests is especially notable because these compounds are highly persistent and lipophilic, properties that favour their accumulation in fur and organic nest material (California Environmental Protection Agency, 2023; Simon-Delso et al., 2015). In addition, fipronil sulfone can persist and accumulate in animal tissues and has been associated with toxic effects in non-target vertebrates (De Marchi et al., 2025; Zhao et al., 2005). Their occurrence despite restrictions on agricultural use within the European Union (European Commission, 2013) further indicates that veterinary applications constitute an important pathway of environmental contamination.

The detection of acetamiprid (a widespread neonicotinoid insecticide) exclusively in urban nests suggests that multiple contamination pathways may operate simultaneously in urban environments. As the only neonicotinoid currently approved for outdoor use in the European Union, acetamiprid remains widely used in agriculture and horticulture across Europe, including domestic gardening in Hungary, where its application against common plant pests is widespread (Varga-Szilay & Pozsgai, 2022). Its presence in nest-lining material therefore likely reflects inputs beyond veterinary treatments, including pesticide use in domestic gardens and urban vegetation management, such as treatments of ornamental horse chestnut (*Aesculus hippocastanum*) trees against key pests (Percival et al., 2012), common in Hungarian cities. This pattern underscores the complexity of urban contaminant mixtures and indicates that birds breeding in cities are exposed to a combination of veterinary and non-veterinary insecticide sources.

Insecticide residues were also detected in nests from the Natura 2000 forest site, indicating that veterinary insecticides can disperse beyond densely populated areas and reach semi-natural and protected ecosystems, albeit at lower concentrations than in urban habitats. Recreational activities likely contribute to this process, as forest trails are frequently used by dog owners (Arnberger, 2006), while hunting dogs may represent an additional source of contaminated fur. Livestock treated with ectoparasiticides, such as cattle and sheep, may further contribute to environmental contamination (Wells & Collins, 2022). Together, these findings demonstrate that protected areas are not isolated from veterinary insecticides and may be subject to chemical inputs arising from a range of human activities, from recreational use to agricultural and livestock management.

Contrary to the first part of our second hypothesis (**H2**), insecticide contamination was lower in second broods than in first broods, likely reflecting a combination of non-exclusive processes. Residues on fur and other nest materials may degrade over time through photodegradation, microbial activity, or weathering, and may also be reduced by rainfall-driven leaching or wash-off, although the relative importance of these processes is likely to vary among compounds. Seasonal variation in companion animal treatment practices may additionally influence the environmental availability of contaminated fur throughout the breeding season.

Our results provide empirical support for the hypothesis proposed by Tassin–de–Montaigu et al. (2026), who suggested that insecticide concentrations in nest materials may decline over the course of the breeding period. By sampling the same nests twice – once during the nestling stage and again after fledging – we were able to directly quantify within-nest temporal variation. Supporting the second part of **H2**, insecticide concentrations decreased between the two sampling points, indicates that residues are dynamic components of nests, with potential implications for exposure during chick development. However, within-nest insecticide concentrations may also be influenced by ongoing nest construction and maintenance, as females are known to add new lining material (e.g. animal fur) during nest renovation (Harnist et al., 2020).

Tassin–de–Montaigu et al. (2026) reported strong positive associations between insecticide contamination in nests and residues detected in unhatched eggs and dead nestlings. In contrast, we found no significant relationships between insecticide exposure and hatching success, nestling mortality, or nestling body size parameters. Our results rather align with Tank et al. (2026), who similarly reported no associations between pesticide concentrations and biometric traits such as body mass and wing length in five common bird species. These findings are notable given that reproductive and morphological parameters were recorded using standardised protocols throughout the breeding period. Collectively, these findings do not support our third hypothesis (**H3**), suggesting that insecticide contamination in nest-lining material does not necessarily translate into measurable effects on the reproductive and morphological endpoints assessed here.

The absence of detectable effects may partly reflect limited statistical power to resolve subtle or sublethal impacts, as well as the fact that the endpoints measured in this study may not capture more complex biological consequences of chronic low-dose exposure. Indeed, experimental and toxicological evidence indicates that compounds such as fipronil and imidacloprid can affect neurological, endocrine, behavioural, and reproductive functions in birds even at low concentrations (Pisa et al., 2021). As a result, effects may manifest through pathways not addressed here, including altered parental care, immune function, or post-fledging survival.

An additional, non-mutually exclusive explanation involves potential trade-offs between the direct toxic effects of insecticides and their indirect ecological effects via reduced ectoparasite loads in nests, as suggested in previous studies (e.g. Suárez-Rodríguez et al., 2013). If these compounds simultaneously reduce nest parasite pressure, potential benefits for chick condition or survival may partially offset sublethal toxic effects. Disentangling these opposing pathways will require studies that jointly quantify parasite communities and nestling development beside insecticide presence, as such compensatory dynamics may be most evident under varying baseline parasite pressures.

Brood type itself significantly affected some reproductive parameters, particularly nestling mortality and tarsus length, with second broods generally performing worse than first broods. Seasonal declines in reproductive performance are common in passerines, and also in our urban Great Tit population (Sinkovics et al., 2023), and are often associated with reduced food availability, poorer prey quality, or increased parasite pressure later in the breeding season (Verhulst et al., 1995). These strong seasonal effects may partly mask more subtle influences of insecticide exposure.

Future perspectives should focus on contamination patterns across urban gradients, as different land-use types likely differ in their contribution to environmental insecticide exposure. Residential areas with high densities of companion animals and frequent dog-walking activity may represent hotspots (Arnberger, 2006; Preston-Allen et al., 2023), while commercial or administrative districts may experience lower inputs. Suburban and peri-urban green spaces, as well as urban parks and recreational forests, may act as important interfaces between wildlife and anthropogenic chemical exposure and therefore warrant targeted investigation.

Future studies should also assess potential effects on adult birds, as exposure is not limited to nestlings. In small-bodied species such as the Great Tit, females may experience higher exposure than males due to their exclusive role in incubation and their primary responsibility for collecting and arranging nest-lining material (Mainwaring, 2017). This risk may be further amplified in urban populations, where females have been reported to spend more time incubating than their forest counterparts (Hope et al., 2022), potentially increasing the duration of contact with contaminated nest material and pointing to the need to investigate possible sex-specific effects.

As evidence for the presence of veterinary insecticides in terrestrial environments increases (e.g. Perkins et al., 2025; Tank et al., 2026), there is also a need to better understand their ecological and physiological relevance using more sensitive approaches. Beyond single-point reproductive measures, future studies should incorporate finer-scale developmental metrics, such as growth trajectories, as well as physiological indicators, including stress-related hormones, to better capture potential sublethal effects in wild bird populations.

## 5. Conclusion

Our study shows that veterinary ectoparasiticides are present in Great Tit nests not only in urban habitats but also in a Natura 2000 forest, suggesting that companion animal treatments represent a previously under-recognised pathway of pesticide exposure in wild birds. These findings indicate transfer from treated companion animals to avian breeding microhabitats via contaminated fur incorporated into nest-lining materials, with consistently higher contamination levels in urban nests, suggesting increased exposure pressure in densely populated human-modified landscapes.

Although we found no clear associations between nest contamination and the measured reproductive parameters, the occurrence of multiple neurotoxic compounds in both habitat types raises ecological concerns. Several of the detected substances are restricted in agricultural contexts within the European Union due to risks to non-target organisms, yet remain widely applied in veterinary contexts and are largely absent from current environmental monitoring frameworks.

Future research should prioritise the investigation of sublethal and long-term effects on wild birds, including impacts on physiology, behaviour, immune function, and post-fledging survival. Comparative studies across urbanisation gradients and other human-influenced green spaces will be essential to identify exposure hotspots and clarify dispersal pathways. Overall, our results highlight the need to incorporate veterinary ectoparasiticides into broader ecological risk assessments of pesticide exposure in wildlife.

### PERMITS

All necessary permits for monitoring bird development, nest surveillance, and work conducted in protected areas are held by the research team (VE-09Z/03454-8/2018 for working in protected areas and PE/EA/786-7/2018 for observing and handling protected species).

## CRediT authorship contribution statement

**Zsófia Varga-Szilay**: Conceptualization, Methodology, Software, Validation, Formal analysis (with advices from GS and IP), Investigation, Resources, Data Curation, Writing - Original Draft, Writing – Review & Editing, Visualization, Project administration, Funding acquisition. **Olivér Csíkvári**: Methodology, Resources, Writing – Original Draft (LC-MS/MS analysis), Writing – Review & Editing, Funding acquisition. **Olga Ulbert**: Methodology, Writing – Original Draft (GC-MS analysis), Writing – Review & Editing. **Ivett Pipoly**: Conceptualization, Methodology, Investigation (avian fieldwork), Writing – Original draft, Writing – Review & Editing. **Gábor Seress**: Conceptualization, Methodology, Investigation (avian fieldwork), Data Curation, Writing – Original draft, Writing – Review & Editing, Funding acquisition. All authors have read and agreed to the published version of the manuscript.

## Code availability

The underlying computer code is available in the GitHub repository https://github.com/zsvargaszilay/veterinary_insecticides_in_bird_nests

## Acknowledgement

We, the authors, thank Krisztián Horváth for solving logistical problems and providing the infrastructural support required for GC-MS analyses. We thank the three field assistants, Viola Nagy, Máté Havasi and András Zámbó, for their help in collecting the nest-lining materials. We also thank Renáta Unger for her assistance with the laboratory work on permethrin measurements and Barnabás Gere for his assistance with sample preparation. We are grateful to Gábor Pozsgai for his ideas during the early stages of the project and for his constructive advice throughout the development of the manuscript. We would also like to thank Zápor and Zazie, the two dogs who provided blank fur matrices for analysis.

