## Supplementary Material for "Veterinary insecticides in wild bird nests: emerging contaminants in urban and protected forest habitats"

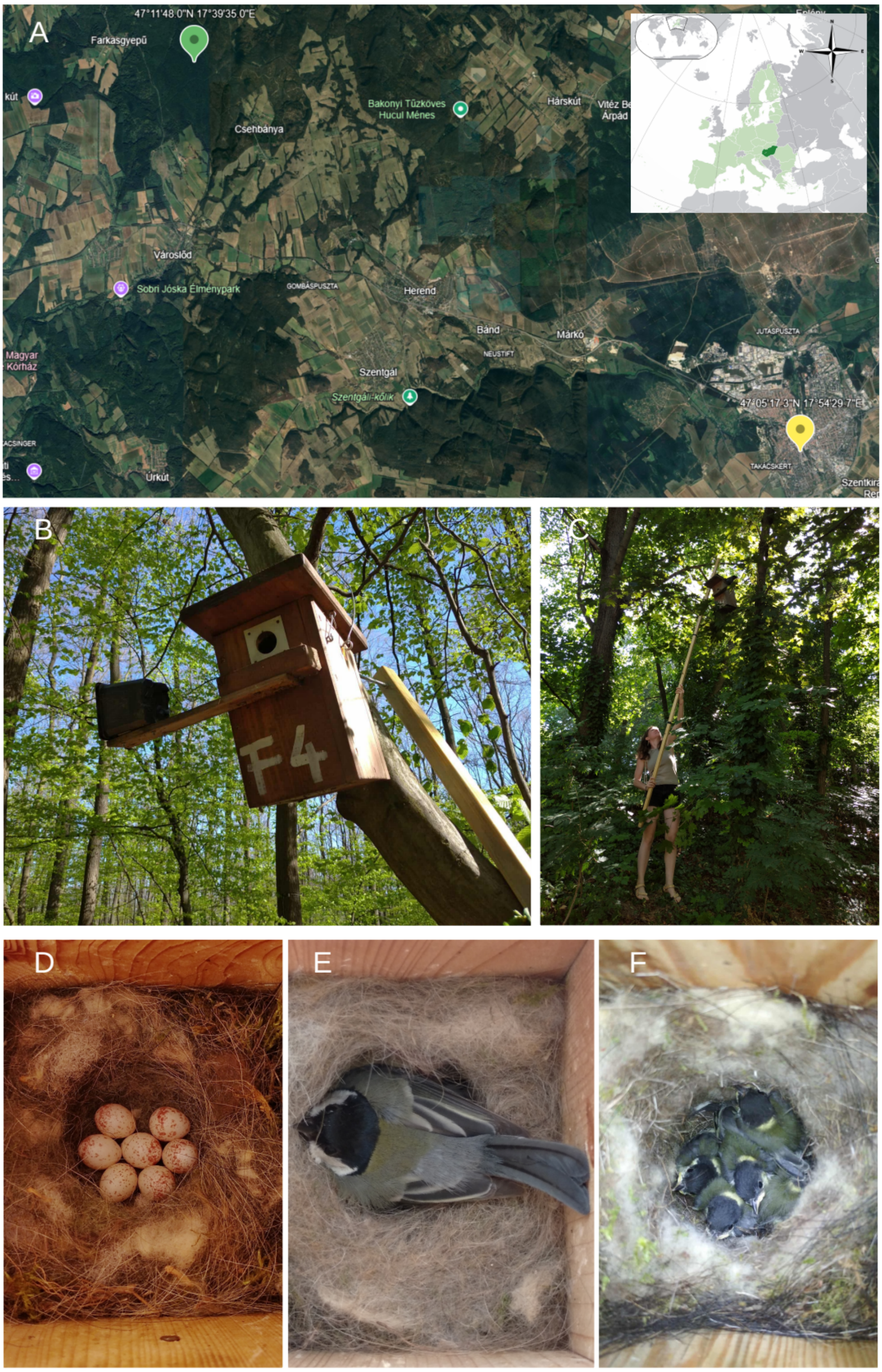

**Supplementary Material Figure 1: Study sites, nest-box system, and breeding stages of Great Tits (*Parus major*).** (A) Location of the two study sites in western Hungary: an urban population in the city of Veszprém (47°05′17.29″N, 17°54′29.66″E), where nest boxes were distributed along an urbanisation gradient, and a forest population near Farkasgyepű in a protected Natura 2000 forest (47°11′48″N, 17°39′35″E). (B) Nest-box in the forest habitat. (C) Nest-box monitoring. (D) Lined nest containing eggs. (E) Female incubating eggs inside the nest box. (F) Late-stage nestlings shortly before fledging. All photographs (B-F) were taken by Zsófia Varga-Szilay.

**Supplementary Material Table 1:** Gradient program for the separation of dinotefuran, fipronil, fipronil sulfone, imidacloprid, and nitenpyram.

| Time [min] | Flow rate [mL/min] | %A | %B |
|---|---|---|---|
| 0.000 | 0.700 | 100.000 | 0.000 |
| 0.120 | 0.700 | 100.000 | 0.000 |
| 0.150 | 0.700 | 100.000 | 0.000 |
| 0.500 | 0.700 | 5.000 | 95.000 |
| 0.550 | 0.700 | 5.000 | 95.000 |
| 0.690 | 0.700 | 5.000 | 95.000 |
| 0.990 | 0.700 | 5.000 | 95.000 |
| 1.000 | 0.700 | 100.000 | 0.000 |
| 1.400 | 0.700 | 100.000 | 0.000 |

**Supplementary Material Table 2:** Optimised Multiple Reaction Monitoring parameters of analytes.

| Analyte | Q1 m/z | Q3 m/z | DP | EP | CE | CXP | Polarity |
|---|---|---|---|---|---|---|---|
| Dinotefuran | 158.35 | 101.92 | 80 | 4 | 20 | 8 | Positive |
| Acetamiprid | 223.1 | 89.87 | 80 | 2 | 45 | 7 | Positive |
| Nitenpyram | 271.38 | 224.98 | 70 | 2 | 15 | 7 | Positive |
| Fipronil | 434.64 | 329.78 | -60 | -10 | -20 | -11 | Negative |
| Fipronil sulfone | 450.58 | 414.76 | -70 | -10 | -20 | -11 | Negative |
| Imidacloprid | 255.95 | 208.99 | 70 | 10 | 20 | 11 | Positive |

**Supplementary Material Table 3:**Lower Limits of Quantitation for analytes quantified by LC-MS/MS.

| Analyte | LLOQ [ng/sample] |
|---|---|
| Dinotefuran | 0.414 |
| Acetamiprid | 0.080 |
| Nitenpyram | 0.200 |
| Fipronil | 0.071 |
| Fipronil sulfone | 0.065 |
| Imidacloprid | 0.193 |

**Supplementary Material Table 4:** GC-MS parameters used for the analysis of sample extracts.

| | |
|---|---|
| Instrument: Shimadzu GCMS-QP2010 SE | |
| Analytical column: ZB-5MS plus (30 m, 0.25 mm, 0.25 µm) | |
| Injector temperature | 300ºC |
| Injection mode | Splitless |
| Volume of injection | 1 µL |
| Column flow | 0.99 mL/min (He) |
| Initial temperature | 60ºC |
| Oven program | 60-310ºC to 20ºC/min |
| Final temperature | 300ºC (3 min) |
| Interphase temperature | 280ºC |
| Ion source temperature | 250ºC |
| Solvent delay time | 2.5 min |
| MS program and monitored isotopes | 2.51-11.00 min: scan (event time: 0.3 s), with detection of 35-350 m/z isotopes<br>11.01-14.00 min SIM (event time: 0.3 s) for detection of 183, 188, 184, 115, 82 m/z isotopes |
